# The spatiotemporal organization of experience dictates hippocampal involvement in primary visual cortical plasticity

**DOI:** 10.1101/2021.03.01.433430

**Authors:** Peter S. B. Finnie, Robert W. Komorowski, Mark F. Bear

**Affiliations:** The Picower Institute for Learning and Memory, Department of Brain and Cognitive Sciences, Massachusetts Institute of Technology, Cambridge, US 02139

**Keywords:** primary visual cortex, hippocampus, synaptic plasticity, long-term memory, systems consolidation

## Abstract

The hippocampus and neocortex are theorized to be crucial partners in the formation of long-term memories. Here, we assess hippocampal involvement in two related forms of experience-dependent plasticity in the primary visual cortex (V1) of mice. Like control animals, those with hippocampal lesions exhibit potentiation of visually evoked potentials following passive daily exposure to a phase reversing oriented grating stimulus, which is accompanied by long-term habituation of a reflexive behavioral response. Thus, low-level recognition memory is formed independently of the hippocampus. However, response potentiation resulting from daily exposure to a fixed sequence of four oriented gratings is severely impaired in mice with hippocampal damage. A feature of sequence plasticity in V1 of controls, but absent in lesioned mice, is generation of predictive responses to an anticipated stimulus element when it is withheld or delayed. Thus, hippocampus is involved in encoding temporally structured experience, even in primary sensory cortex.

## Introduction

Distinctions between hippocampus-dependent and -independent memories may include the locus of information storage, the types of information stored, and the mechanism(s) of encoding and consolidation (*1–5*). In recent years it has been established that mouse primary visual cortex (V1) is a storage site for several types of memory historically considered to be the domain of “higher” brain regions (*6*), offering a new opportunity to understand the nature of hippocampus-dependent encoding in neocortex. The advantages of studying mouse V1 are that [1] sensory experience can be precisely controlled, [2] the resulting modifications can occur prior to binocular integration and depend on mechanisms local to V1, pinpointing this as a critical locus of storage, and [3] the experience-dependent plasticity is reported by robust changes in visual-evoked potentials (VEPs) (*7*). However, it remains to be established if any of these experience-dependent modifications of V1 depend on hippocampus, and if they do, what distinguishes them.

In the current study, we have compared the hippocampal dependence of two types of V1 plasticity that have in common potentiation of the VEP. Both forms of plasticity are triggered by brief daily exposure of awake head-fixed mice to carefully controlled visual stimuli. The first protocol consists of exposure to a phase-reversing, oriented grating stimulus, which elicits stimulus-selective response plasticity (SRP) expressed as an increase in VEP magnitude recorded in V1 (*8*). This response modification possesses attributes consistent with perceptual learning, including gradual emergence in the hours following experience, persistence over weeks, and exquisite stimulus selectivity (*9, 10*). SRP is accompanied by habituation of an innate behavioral response to presentation of the visual grating stimulus, indicating formation of long-term recognition memory (*11*). Both the electrophysiological and behavioral measures of memory formation are disrupted by local manipulations of N-methyl-d-aspartate (NMDA)-receptors and inhibitory neurotransmission within V1, suggesting a common mechanistic basis in neocortex (*7, 12*). The second visual stimulation protocol comprises repeated daily exposure to four distinct grating orientations, arranged in a consistent temporal order. Like SRP, response potentiation emerges gradually in the hours after stimulation, is highly selective for stimulus properties present during training, occurs prior to binocular integration, and is reliant on plasticity mechanisms local to V1 (*13*). However, unlike SRP, visual sequence plasticity depends on local activation of muscarinic cholinergic receptors but does not require NMDA-receptors in V1. Thus, SRP and sequence potentiation are similarly expressed by VEPs in V1, but are driven by qualitatively different types of experience and depend on distinct mechanisms.

In the current study we demonstrate that these two similar forms of V1 response potentiation also have dissociable reliance on the hippocampus. In mice with dorsal and ventral hippocampal damage, both SRP and long-term behavioral habituation are largely normal, but V1 potentiation elicited by visual sequence exposure is virtually absent. Control mice, familiarized to a consistent sequential pattern of visual stimulation, produce anticipatory V1 responses even when an element is omitted or delayed. Hippocampal damage eliminates this generative V1 response, providing evidence that interactions between these regions are necessary for spatiotemporal prediction (*14*). Moreover, following daily exposure to a series of gratings, hippocampectomized mice undergo far less potentiation not just to the familiar sequential arrangement, but also when the stimuli are presented in reverse order, suggesting a failure to encode or ‘recognize’ each constituent grating orientation. These findings suggest that the hippocampus contributes to long-term plasticity in primary sensory cortex under circumstances when discrete stimuli are arranged in a predictable temporal order.

## Results

### Long-term visual recognition does not require the hippocampus

To assess hippocampal involvement in low-level visual recognition memory, we recorded extracellular local field activity from layer 4 (L4) of binocular V1 along with concomitant forepaw movement while head-restrained mice viewed an oriented grating stimulus over multiple daily sessions. We use the term “vidget” to describe the transient increase in forepaw movement when the visual stimulus transitions from gray to grating (*11*). To eliminate hippocampal influences, we produced permanent bilateral lesions prior to experimentation via NMDA nanoinjections into the hippocampus (*15*). **Figures 1A-B** and **S1** display representative examples of bilateral hippocampal lesions that met inclusion criteria, alongside coronal sections from the brains of littermate control mice that received sham lesions. Post-mortem histology determined that the area of gross residual hippocampal tissue following lesioning was 40.87 ± 10.22% that of sham mice (mean ± SEM = 100 ± 4.34%), a statistically significant difference (Mann-Whitney U = 9, p = 0.0001).

**Figure 1.**
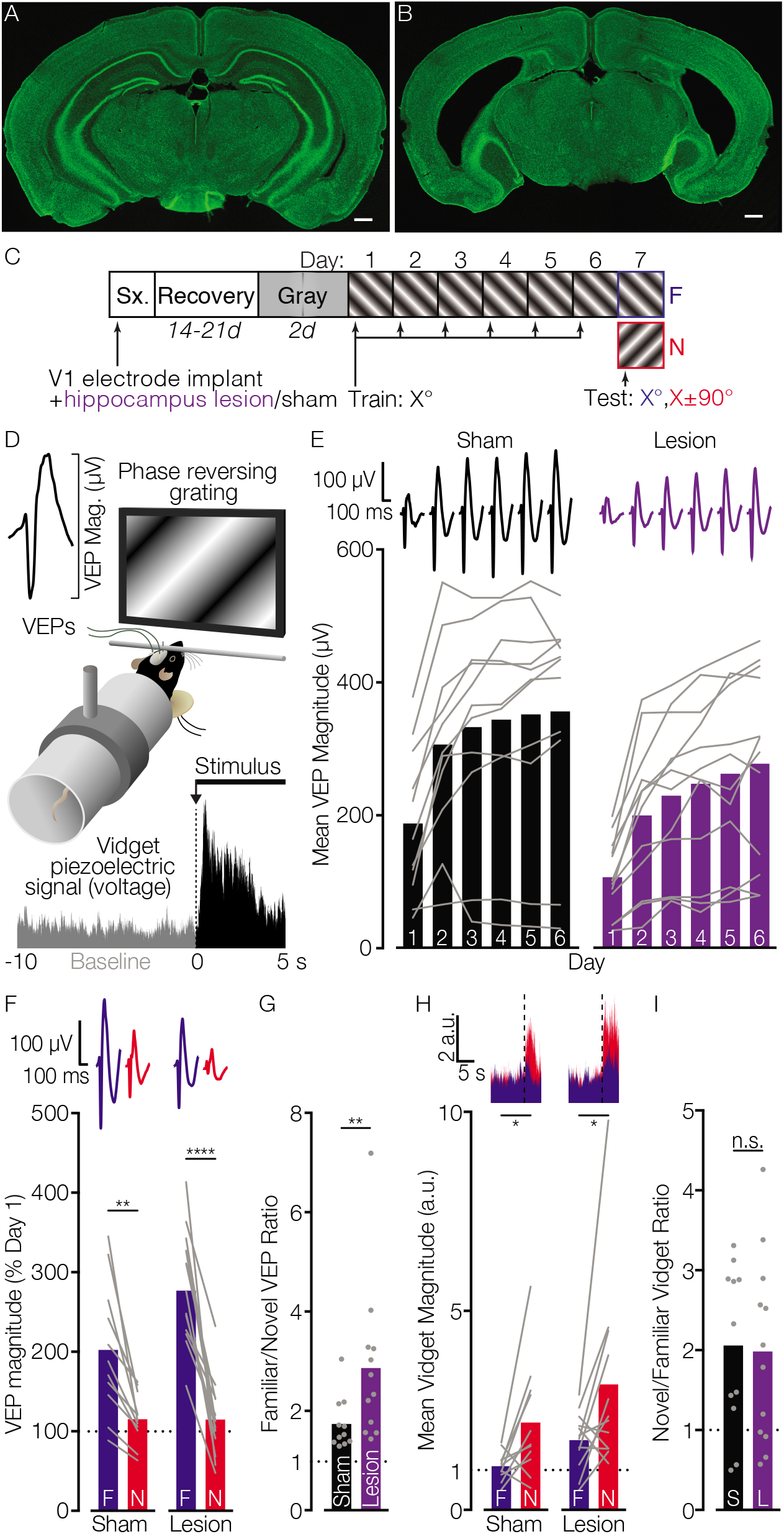
Long-term visual recognition memory does not require the hippocampus. Representative coronal brain sections processed via NeuN immunofluorescence from mice in (**A**) sham control and (**B**) hippocampal lesion groups. Green, NeuN. Scale bar represents 500 μm. Diagrams depict the **(C)** experimental protocol and **(D)** apparatus. **E)** Mean VEP magnitudes plotted for each training day (days inset along x-axis, mean daily voltage traces plotted at top, and lines depict mean magnitude for each mouse). In both groups VEPs potentiated across days of exposure to a grating stimulus (two-way RM ANOVA with Greenhouse-Geisser correction, effect of day [*F*_1.859,39.04_ = 50.96, *p* < 0.0001]; Sidak’s posthoc tests comparing Days 1 to 2-6, each *p* < 0.0001). However, potentiation did not differ between Sham and Lesion groups (neither a main effect of group [*F*_1,21_ = 2.89, *p* = 0.1] nor group by day interaction [*F*_1.859,39.04_ = 0.44, *p* = 0.63]). **F)** VEP magnitudes elicited by familiar (F) and novel (N) stimulus orientations on *Day 7*, normalized to *Day 1*. The difference in familiar versus novel responses was significantly larger in the Lesion relative to Sham group (RM ANOVA group by stimulus interaction; *F*_1,21_ = 6.03, *p* = 0.023; Sidak’s post hoc tests: F vs N, Sham *p* = 0.0017, Lesion *p* < 0.0001). **G)** The ratio of VEP magnitudes elicited by F relative to N was also larger in the Lesion versus Sham group (two-tailed Mann-Whitney *U* = 24, *p* = 0.009), confirming that SRP is exaggerated following hippocampal ablation. **H)** In each group, larger mean behavioral response magnitudes (arbitrary units, a.u.) were elicited by the onset of N versus F stimuli on Day 7 (mean traces superimposed at top; two-tailed Wilcoxon matched-pairs planned comparisons: sham, *Z* = 2.223, *p* = 0.024; lesion, *Z* = 2.118, *p* = 0.034). **I)** The N/F vidget ratios for each group were comparable (unpaired two-tailed *t*-test, *t*_21_ = 0.16, *p* = 0.88). \**p* < 0.05, \*\**p* <0.01, *** p < 0.001, \*\*\*\**p* < 0.0001, n.s. *p* > 0.05. *n*/group: Sham = 11, Lesion = 12.

After lesion and sham control mice acclimated to head-fixation across two daily sessions (while viewing a uniform gray screen), they were exposed to an isoluminant, phase-reversing grating stimulus of a fixed orientation for six consecutive days (**Fig. 1C-D**). On Day 1 (baseline), mean VEP magnitudes were significantly smaller in the lesion (106.1 ± 16.64 μV) relative to sham (187± 33.71 μV) group (two-tailed *t*-test, *t*_21_ = 2.21, *p* = 0.038). The cause(s) of the baseline response magnitude difference remain unidentified, but could include altered electrical volume conduction (*16*), network synchrony (*17–21*), mild diaschisis within the thalamocortical visual circuit (*22*), or shifting V1 electrode placement due to atrophy of the underlying hippocampal tissue. Nevertheless, behavioral responses to the onset of grating stimuli presented on Day 1 were comparable in each group (normalized to pre-stimulus baseline; unpaired *t*-test, *t*_21_ = 1.53, p = 0.14; **Fig. S2**), as was spontaneous movement during interleaved presentations of gray screen between blocks (unpaired *Mann-Whitney U* test; sham: *Z* = 52, *p* = 0.41). Importantly, across six days of exposure to the same oriented grating stimulus, VEP magnitudes potentiated significantly with no detectable differences between groups (**Fig. 1E**). Thus, SRP acquisition was normal in mice with hippocampus lesions. On Day 7, the responses to the familiar stimulus were also compared to a stimulus with a novel orientation. Relative to novel, the VEP magnitude to the familiar stimulus was greater in both groups, but significantly exaggerated in lesioned mice (**Fig. 1F-G**). The enhanced potentiation could be attributable to the smaller baseline VEP magnitudes measured in hippocampectomized mice, or to improved memory (*23–25*). Regardless, an intact hippocampus is evidently not critical for this form of experience-dependent V1 plasticity. Concomitant orientation-selective habituation (OSH) of the vidget was likewise observed in both groups (**Fig. 1H**), which did not differ statistically (**Fig. 1I**). A separate cohort of mice that received lesions targeting only the dorsal hippocampus (dH) likewise acquired and expressed both SRP and OSH at normal levels (**Fig. S3**). Thus, the hippocampus is not required for V1 plasticity underlying long-term visual recognition memory.

To test whether the hippocampus is required for the retention and/or consolidation of SRP over longer time intervals (*26*), a subset of mice reported above also received a second test session 10 days following completion of the 7-day protocol. SRP persisted in both the sham and lesion groups (**Fig. S4**), indicating that plasticity declines little across this retention interval, even when formed in the absence of hippocampus. To verify the functional ablation of hippocampus, we next subjected all mice exposed to the SRP protocol to an object displacement behavioral task known to be sensitive to hippocampal dysfunction (*27, 28*) (**Fig. 2**). In both groups, the exploration of two identical objects in static locations diminished significantly across four sampling sessions. However, during a final test session, mice in the sham condition preferentially explored the object that had been moved to a new spatial location whereas lesioned mice investigated both objects for an equivalent duration. This finding is consistent with bilateral damage to the hippocampus.

**Figure 2.**
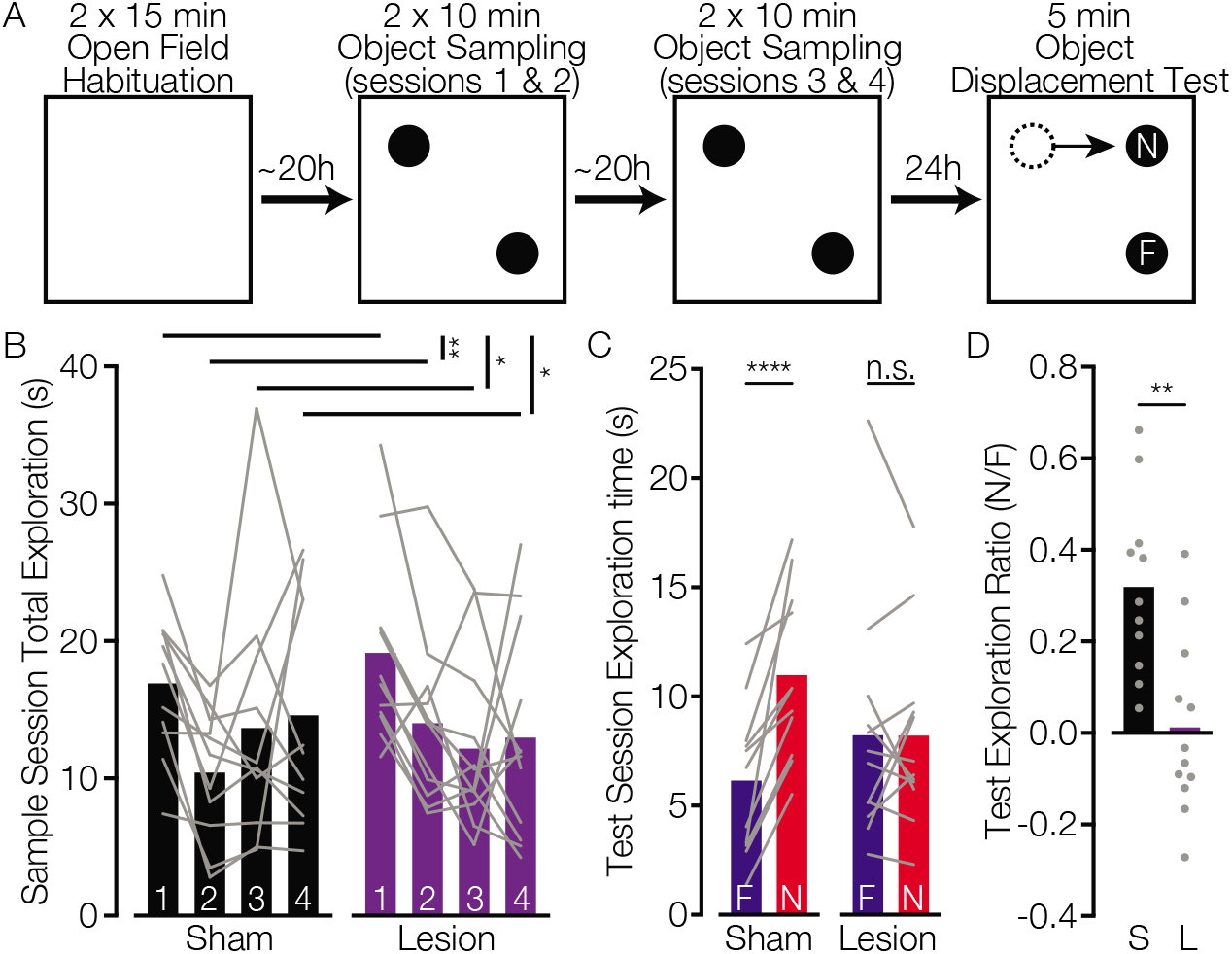
Pre-training hippocampal lesions impair spontaneous exploration of a spatially-displaced object. **A)** Diagram of experimental timeline and apparatus. Square boxes represent overhead views of the open field arena, and filled circles example positions of identical objects during sampling and testing phases. **B)** The average time mice explored two identical objects declined between the first and each subsequent sampling sessions (two-way RM ANOVA with Greenhouse-Geisser correction, main effect of session, *F*_2.071,43.482_ = 4.38, *p* = 0.017; Dunnetts’s posthoc tests comparing Sessions 1 with 2 (*p* = 0.005), 3 (p = 0.013), and 4 (p = 0.049)). There was no difference in exploration between Lesion and Sham groups during the sampling sessions (main effect of Group, F_1,21_ = 0.13, *p* = 0.73; Group by Session interaction, *F*_2.071,43.482_ = 1.13, *p* = 0.34). **C)** On test day the Sham group explored a spatially displaced object for a significantly longer duration (N mean = 10.97±1.17 s) than a static object (F mean = 6.15±1.04 sec; treatment by object interaction: F_1,21_ = 14.03, *p* = 0.001; Sidak’s multiple comparison test, *p* < 0.0001), whereas the lesion group explored both objects equivalently (F = 8.23±1.55 sec; N = 8.21±1.25 s; Sidak’s, *p* > 0.99). **D)** A ratio of exploration times for the N versus F object was also significantly greater for the Sham (0.31 ± 0.05 a.u.) relative to Lesion (0.012 ± 0.06 a.u.) group (two-tailed unpaired *t*_21_ = 3.79, *p* = 0.001). * < 0.05, ** < 0.01, **** < 0.0001. *n*/group: Sham = 11, Lesion = 12.

### Hippocampal damage disrupts potentiation of V1 responses following visual sequence exposure

Hippocampal perturbations can disrupt the ability to learn the spatial and temporal arrangement of previously encountered items (*15, 29, 30*). Thus, primary sensory cortex alone may be sufficient to form simple stimulus representations, but relational memory might additionally require the hippocampus (*28, 31, 32*). To investigate this possibility, we interrogated a form of VEP potentiation that is evoked in V1 when mice are exposed daily to gratings of 4 different orientations in a fixed sequence (referred to here as stimulus *A, B, C*, and *D*; **Fig. 3A**) (*13*). As observed on Day 1 of the SRP protocol, mean baseline VEP magnitudes elicited by the four stimuli in the visual sequence were marginally larger in the sham group relative to the lesion group, however this did not reach statistical significance (two-tailed Welch’s *t*-test, *t*_8.42_ = 0.73, *p* = 0.485). Across consecutive training days, sham control mice exhibited robust potentiation of mean VEP responses elicited by the stimulus sequence, whereas lesioned mice exhibited no significant change in VEP magnitude (**Fig. 3B**). To dissociate sequence-from stimulus-selective potentiation, on Day 5 mice were exposed to the same set of oriented gratings, presented in both forward (*ABCD*) and backward (*DCBA*) order. As the first stimulus in the familiar sequence (‘*A*’) appears to predictively cue neural response modulation to gratings presented thereafter, potentiation is typically most pronounced for the two middle elements (*13*). Thus, to assess the effect of hippocampal lesions on V1 plasticity evoked by spatiotemporal patterns we compared VEPs elicited by elements *B* and *C* in the forward and backward sequences, after normalizing to mean Day 1 magnitude for each group (**Fig. 3D**). Only sham control mice exhibited significant sequence-specific potentiation, which was absent in the lesion group (**Fig. 3E**). Surprisingly, the normalized response magnitude evoked by the reverse sequence (*CB*) was also significantly larger in the Sham group than the Lesion group. This finding suggests that direction-invariant potentiation of VEPs, putatively elicited by familiarity with each discrete oriented stimulus, is absent in hippocampectomized mice. This stands in stark contrast to the supranormal SRP observed in lesioned mice (**Fig. 1**).

**Figure 3.**
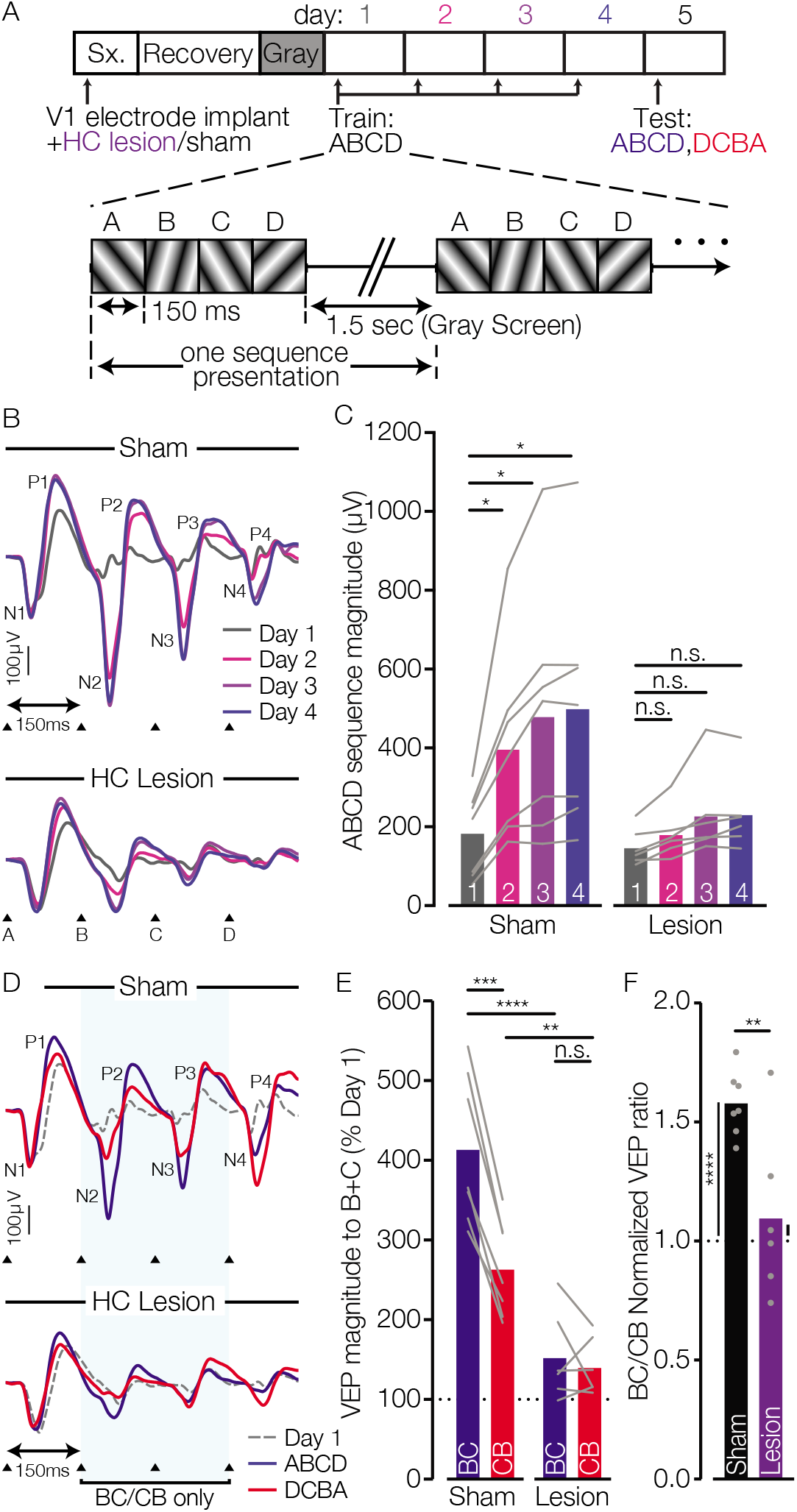
The hippocampus is required for V1 response potentiation evoked by a sequence of visual stimuli. **A)** Schematic diagram of experimental time-course and daily visual stimulation protocol. **B)** Average VEP waveforms elicited by visual sequence *ABCD* for each group on *days 1-4*. Labeled arrows denote the onset latency of sequence elements A, B, C, and D. N1-4 and P1-4 labels refer, respectively, to the four peak negative- and positive-going deflections of the V1 local field potential following each stimulus onset. **C)** Potentiation of average VEP magnitude to the familiar sequence across *Days 1-4* was impaired in mice with hippocampal relative to sham lesions (two-way RM ANOVA, day by group interaction, *F*_3, 33_ = 6.20, *p* = 0.0018). Within-subject potentiation was significant for the Sham (Sidak’s posthoc comparisons of *Days 1* versus *2-4*: *p* = 0.024, 0.027, and 0.022, respectively) but not Lesion group (*Day 2*, *p* = 0.072; *Day 3, p* = 0.11; *Day 4, p* = 0.061). **D)** Average VEP waveforms elicited by sequence *ABCD* and the reverse *DCBA* during the *Day 5* test session (*Day 1* superimposed for reference). **E)** On *Day 5*, response potentiation in Lesion and Sham groups were significantly different for elements B and C in the *ABCD* versus reverse *DCBA* sequences, normalized to the *day-1* baseline magnitudes for each mouse (two-way RM ANOVA group by stimulus interaction, F_1, 11_= 31.73, *p* = 0.0002). Responses to *BC* were larger in the Sham controls than in the Lesion group (planned Sidak’s comparison, *p* < 0.0001), suggesting the hippocampus is required for sequence-specific potentiation. Furthermore, response magnitude to the forward (*BC*) sequence was only larger than the reverse (*CB*) in the Sham (*p* < 0.0001) but not the Lesion group (*p* = 0.75). Mean response to the reverse *CB* sequence was also significantly larger for the Sham compared to Lesion group (*p* = 0.0057), indicative of an overall difference in sequence-independent potentiation. Subsequent planned one-sample *t*-tests with Bonferroni correction revealed significant potentiation of responses to both *BC* and *CB* over the *Day-1* baseline only in the Sham (*p* = 0.0004 and 0.0016, respectively) but not the Lesion group (*p* = 0.26 and 0.15, respectively). **F)** The familiar/novel ratio on *Day 5* was also significantly larger in the Sham versus Lesion group (Welch’s two-tailed *t*-test, *t*_6.316_= 3.162, *p* = 0.018). Furthermore, planned one-sample *t*-tests confirmed that sequence-specific potentiation was only evident in the sham (*p* < 0.0001) but not lesion (*p* = 0.51) group. Bars represent group means and points/lines are mean values for each individual mouse. \**p* < 0.05, \*\**p* <0.01, *** p < 0.001, \*\*\*\**p* < 0.0001, n.s. *p* > 0.05. *n*/group: Sham = 7, Lesion = 6.

To directly confirm that hippocampectomy has differential effects on sequence learning versus SRP, we performed a Kruskal-Wallis test on the Familiar/Novel ratios for each group. These groups differed statistically, χ^2^(3) = 16.79, p = 0.0008, and planned contrasts with Dunn’s correction for multiple comparisons identified a significant difference in Familiar/Novel ratios for SRP versus sequence learning in hippocampectomized (p < 0.0001) but not sham animals (>0.99). Thus, these two forms of V1 plasticity have dissociable reliance on the hippocampus.

### The hippocampus is required to generate anticipatory V1 responses

Our findings are consistent with the hypothesis that the hippocampus contributes to temporal patterning (*33, 34*)—modulating V1 response magnitude when cued by the first element in a familiar visual sequence. Previously it has been shown that even when sequence elements are omitted, V1 displays sequential reactivation of spatiotemporal neural response patterns (*13, 35*). It has been argued that plasticity within local intracortical circuits may be sufficient for such temporally organized memory (*36*), leaving open the question of hippocampal involvement. To test this directly, we assessed responses to omitted stimuli using two different approaches.

First, during the Day 5 test session we included a modified version of the familiar forward ABCD sequence in which each grating stimulus was held on screen for 300 ms (twice the standard 150 ms duration). We reasoned that when cued with stimulus A, mice would expect the onset of stimulus B approximately 150 ms later and display an anticipatory response even when no visual transition had occurred. Indeed, after 4 prior days of exposure to the ABCD sequence, the late N2-P2 components of the field potential that followed presentation of stimulus A were exaggerated in the sham group, suggestive of an anticipatory response at the expected latency (**Fig. 4A**). In the lesion group, this response modulation was far less pronounced. To determine if this exaggerated response was merely a consequence of the slightly larger VEPs in the sham group, we normalized the N2-P2 magnitude to the preceding N1-P1 components evoked by stimulus A (which should exhibit minimal predictive modulation). Even after normalization, sham control mice displayed significantly larger anticipatory N2-P2 responses, prior to the onset of delayed stimulus B (**Fig. 4C**). An alternative possibility is that the potentiated N2-P2 response observed in sham mice does not reflect an anticipatory response, but rather the late components of the VEP when not interrupted 150 ms later by another stimulus. If this were the case, mice with hippocampal damage might simply possess impaired long-latency VEP responses. To investigate this possibility, we quantified the trough-to-peak N2-P2 magnitude elicited by the familiar stimulus on Day 7 of the SRP protocol (re-analyzed from **Fig. 1**), in which there should be no expectation of stimulus B at the typical latency of the N2-P2 components. As expected, we found that N2-P2 response magnitudes (again normalized to the preceding N1-P1 VEPs) were now comparable in sham and lesion groups (**Fig. 4B-C**), indicating that hippocampal damage does not impair the generation of long-latency evoked responses within the SRP protocol.

**Figure 4.**
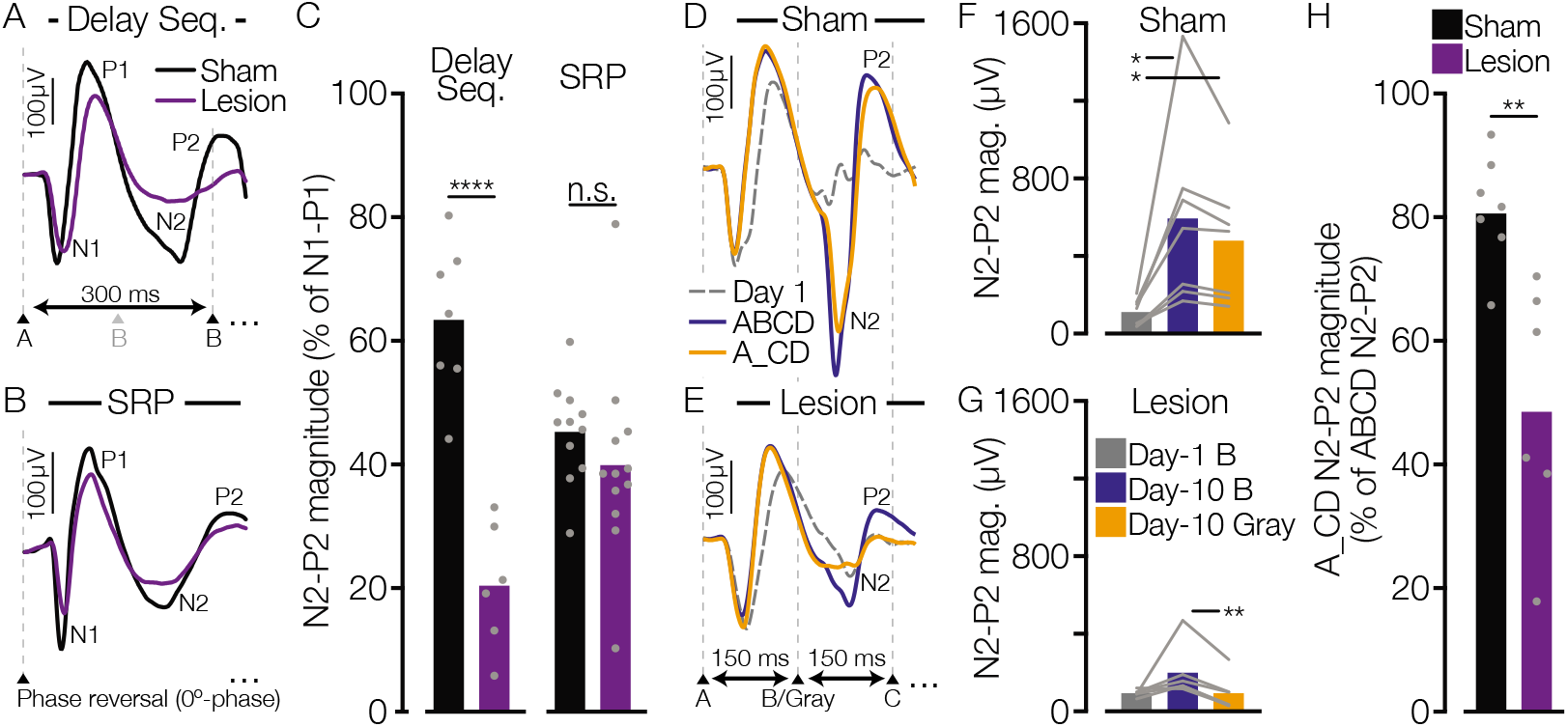
Hippocampal lesions impair the generation of anticipatory responses in V1. **A)** In addition to presentation of the forward and reverse sequences, on Day 5 of the visual sequence protocol we included interleaved blocks of the forward ABCD sequence in which each element was displayed for twice the standard duration (300 ms instead of 150 ms). Traces depict truncated mean VEP waveforms recorded in Sham and Lesion groups following sequence onset, encompassing both early (N1, P1) and late (N2, P2) VEP components elicited by stimulus A. Hashed horizontal lines indicate the transition points between sequence elements (denoted by labeled arrows at panel bottom, with grayed ‘B’ indicating the standard stimulus onset timepoint). **B)** Truncated mean VEP waveforms elicited in Sham and Lesion groups by the familiar grating stimulus on Day 7 of the SRP protocol (re-plotted from Fig. 2). **C)** Graph displays mean N2-P2 responses elicited by stimulus A in the slowed ABCD pattern on Day 5 of the visual sequence protocol (Delay Seq.) and by the familiar grating orientation on Day 7 of the SRP protocol. To account for baseline VEP differences between groups and protocols, the trough-to-peak magnitude of the N2-P2 response was normalized to the average magnitude of P1-N1 for each mouse ([P2 – N2]/[P1 – N1] x 100). Bars represent group means, with individual data-points depicting the mean response magnitudes for each mouse. A significant group-by-protocol interaction (two-way between-groups ANOVA, *F*_1,32_ = 19.31, *p* = 0.0001) is driven by a larger P2-N2 response in sham controls than lesioned mice exposed to the visual sequence protocol (Tukey’s pairwise comparison, p < 0.0001), but not the SRP protocol (p = 0.51). **D-E)** Truncated mean VEP waveforms recorded from Sham control and Lesion groups in response to the ABCD visual sequence on experimental days 1 and 10, as well as A_CD on day 10 (wherein gray screen is substituted for grating stimulus B). **F-G)** Graphs plot mean N2-P2 VEP magnitudes for the Sham and Lesion groups elicited by the 2^nd^ element in each visual sequence (grating orientation B or gray screen). A significant group-by-sequence interaction effect (two-way RM ANOVA with Greenhouse-Geisser correction, *F*_2, 22_ = 5.553, *p* = 0.011) was driven by robust VEP potentiation in the Sham group elicited by stimulus B on Day 10 versus Day 1 (Tukey’s pairwise comparisons, p = 0.0499) and by gray screen on Day 10 versus stimulus B on Day 1 (p = 0.028), but not between stimulus B and gray screen on Day 10 (*p* = 0.198). Conversely, responses in the Lesion group to stimulus B on Day 10 are exaggerated when compared to those elicited by gray screen (Tukey’s Day 10 ABCD vs A_CD comparison, *p* = 0.0079), despite a lack of response potentiation to stimulus B between Days 1 and 10 (*p* = 0.244). **H)** To directly compare generative anticipatory responses between the two groups, we analyzed the N2-P2 responses in the A_CD condition (as a percentage of the ABCD condition on Day 10). The anticipatory VEP during the omission of stimulus B was significantly larger in the Sham than the Lesion group (Welch’s two-tailed t-test, *t*_7.025_ = 4.552, *p* = 0.0026). \**p* < 0.05, \*\**p* < 0.01, \*\*\*\**p* < 0.0001, n.s. *p* > 0.05. *n*/group: Seq., Sham = 7, Lesion = 6; SRP, Sham = 11, Lesion = 12.

Next, to further assess anticipatory response generation in V1, on experimental day 10 we again presented the familiar sequence ABCD along with a variant in which gray screen was substituted for stimulus B (A_CD) (*13*). In sham control mice we confirmed that presentation of stimulus A led to generation of large amplitude N2-P2 responses at the time when stimulus B was anticipated, even when the oriented grating was omitted (**Fig. 4D-G**). In contrast, the lesion group displayed neither significant response potentiation when comparing the VEPs elicited by stimulus B on Days 1 and 10, nor an anticipatory response during gray screen when cued by stimulus A. These data suggest a failure to “fill-in” the predictable oriented grating following hippocampal damage. As a direct comparison of predictive response modulation between groups, mice with hippocampal lesions likewise displayed significantly smaller VEPs to gray screen when normalized to their response to stimulus B on Day 10 (**Fig. 4H**). Together, these findings are consistent with a specific role for hippocampus in predictive response generation during exposure to familiar spatiotemporal patterns of visual stimulation.

Thus far we have inferred a role for hippocampus in spatiotemporal encoding because lesions impaired sequence-evoked plasticity, but not SRP. In part, this interpretation rests on the assumption that SRP principally reflects familiarity with the spatial characteristics of an oriented grating stimulus (*11*), but not the temporal properties of training. However, like the sequence protocol, the visual stimulation pattern routinely used to induce SRP also possesses a stereotyped temporal structure (typically 2 Hz phase reversals). Thus, SRP might likewise involve spatiotemporal prediction. Consistent with this idea, SRP expression possesses emergent properties, manifesting not at the onset of a train of familiar visual stimuli but rather on subsequent phase reversals (*37–39*). If SRP were revealed to be strongly dependent on temporal characteristics of visual experience (as it is for spatial features (*40*)) this would challenge our interpretation that hippocampal lesions selectively disrupt sequential pattern completion. To clarify whether SRP reflects familiarity with stimulus identity or anticipatory prediction of a temporal pattern, a cohort of neurotypical mice was exposed daily to a static grating stimulus After 6 days of exposure, SRP was tested by presenting the familiar stimulus as well as a novel orientation (in interleaved blocks), each phase-reversing at 2 Hz to evoke time-locked responses (**Fig. S5**). Even though the experience during training did not entail strong synchronous V1 activation triggered by phase reversals, robust VEP potentiation was still observed following passive daily exposure to a static grating stimulus. These results demonstrate that SRP does not depend on a predictive response for each upcoming stimulus.

In an additional experiment we integrated the SRP and visual sequence stimulation patterns into a single protocol. Mice viewed two different pairs of gratings during each daily recording session (one consisting of contrast-inverted stimuli with the same orientation, the other consisting of two stimuli with distinct orientations). Response potentiation was exaggerated for the orientation-shifted relative to the phase-shifted set (**Fig. S6**). This indicates that an expected shift in stimulus orientation confers additional anticipatory response modulation on top of the sustained potentiation triggered by stimulus recognition.

## Discussion

Previous studies established that essential synaptic modifications for both SRP and sequence plasticity reside specifically within V1 (*6, 7*), but the possibility of hippocampal involvement was unknown. We found that SRP and the associated long-term behavioral habituation to stimuli recognized as familiar occur independently of the hippocampus. In sharp contrast, hippocampal ablation severely disrupted potentiation of VEPs elicited by a specific sequence of oriented gratings viewed repeatedly across days. Thus, the hippocampus appears to support some forms of long-term experience-dependent plasticity within V1, contingent on the precise spatiotemporal structure of visual stimulation.

Our data are consistent with the notion that hippocampus is important for relational memory by indexing stimulus representations stored in V1 (*32, 41*). A classical taxonomy of long-term memory systems posits that neocortical plasticity alone can support low-level mnemonic operations like sensory priming and perceptual learning, whereas the hippocampus is additionally required to encode declarative information, including episodic memory (*4*). In nonhuman animals, “episodic-like” memory has often been approximated using spatiotemporal tasks that capture defining phenomenological features (i.e. the ability to recall when and where a specific item has been encountered) (*34, 42*). For instance, rodents with hippocampal damage generally learn the identity of objects encountered as they explore an environment, but not their relative positions in space and/or time (*43–45*) (**Fig. 2**). Our findings collected via V1 electrophysiological recordings largely support this distinction. During SRP, stimulus representations appear to be encoded in such a manner that they are dissociable from their surrounding spatial and temporal context (**Fig. S5**) (*11*). However, when multiple grating stimuli are arranged in a consistent sequential pattern, the resultant V1 potentiation is impaired by hippocampal ablation. As sequence-specific potentiation is eye-specific and requires local plasticity mechanisms in V1 (*13*), a simple model is that sensory neocortex locally stores the identity of low-level stimuli and recruits the hippocampus to preserve the temporal relationships among them (*30*). This is consistent with theories positing that the hippocampus does not form a complete, independent record of sensory experience, but rather encodes temporally structured experience by indexing discrete stimulus features stored in neocortex (*46*). Indeed, anticipatory response modulation within a familiar sequence inherently involves pattern completion (the ability to retrieve a complete record of sensory experience when provided a subset of the original cues), which has been attributed to autoassociative network properties conferred by the anatomical connectivity of the hippocampus (*33, 47, 48*). Consistent with the idea that hippocampus has a special role in binding relational properties in widely distributed neocortical circuits, recent work has shown that encoding of visual sequences occurs broadly, extending beyond V1 to include anterior cingulate cortex (ACC) (*49*). We note, however, that although ACC is one of the major intermediaries connecting hippocampus with V1, unlike hippocampus, it is not required for sequence-specific potentiation of VEPs in V1 (*49*).

Although the cortico-hippocampal index model is intuitively appealing, our findings raise an interesting caveat. An implicit assumption of the index model is that neocortex encodes discrete stimulus representations while the hippocampus is necessary to form associations among them (at least initially following experience). Thus, the theory assumes that the neural representation for a given visual stimulus will be stored in the same way by V1 regardless of the preceding and subsequent sensory input. However, at the mechanistic level, this is clearly not the case. Blocking activity of NMDA-receptors prevents SRP induction but has little effect on sequence-specific potentiation. Conversely, blocking muscarinic cholinergic receptors locally in V1 disrupts sequence-specific potentiation but has little effect on SRP (*13*). Thus, there exist in V1 distinct molecular mechanisms capable of achieving precise orientation-selective stimulus encoding, and these may be differentially recruited by the hippocampus. Distinct encoding mechanisms could explain our puzzling finding that mice with hippocampal lesions show significantly less VEP potentiation than the sham group when stimuli are presented in the reverse sequence on test day (**Fig. 3E**). It appears as though hippocampectomized mice fail to recognize each discrete stimulus orientation as familiar specifically when these had been arranged in a sequential pattern throughout training, even though V1 potentiation and behavioral habituation are both normal in the SRP protocol (**Fig. 1**). Therefore, how the brain stores long-term representations of individual oriented gratings appears to vary depending on their relationship to other sensory stimuli. This conclusion is consistent with observations that neocortical representations of sensory stimuli are strongly influenced by other environmental cues, such as spatial position, and that this modulation requires the hippocampus (*50*).

Limitations on the interpretation of this study are imposed by our use of permanent lesions. As such, we cannot dissociate the specific contributions of the hippocampus in the acquisition, consolidation, and/or expression of visual sequence potentiation. Nevertheless, the current findings address hypotheses on the roles of hippocampus in the storage of information via synaptic plasticity in V1, particularly with regard to consolidation. First, the observation that SRP persists over many days in lesioned animals confirms that not all long-lasting forms of neocortical information storage require hippocampus to be consolidated. Second, although sequence-specific potentiation in V1 is absent in lesioned mice, the properties of this type of plasticity challenge traditional theories of hippocampal involvement in “systems” consolidation. A prevailing view is that rapid plasticity in the hippocampus temporarily stores ongoing experiences until slower intracortical plasticity can gradually strengthen sparse connections among discrete functional ensembles (*47, 51–53*). Available evidence suggests this handoff from hippocampus to neocortex is achieved by spontaneous reactivation of neural activity patterns during offline rest periods (*52, 54–57*). However, if this happens in the case of sequence-specific potentiation in V1, it must occur over a far more rapid time-course than is traditionally assumed. A single exposure to novel visual sequences over ~10 minutes leads to sizeable and stable VEP potentiation that is nearly asymptotic 1 day later (**Fig. 3C**). Furthermore, the offline reactivation of visual sequences stored in hippocampus alone seems unlikely to drive sequence-specific response potentiation in V1, as it is difficult to imagine how this could preserve eye-specificity (*13*).

In conclusion, we demonstrate that long-term neocortical plasticity emerging in the hours following experience can—for some forms of passive sensory stimulation—depend on the hippocampus. Our results are broadly consistent with classical divisions of mnemonic function across brain regions, with V1 storing local representations of low-level visual stimuli and the hippocampus participating in the encoding of higher-order relationships among multiple items. When cued by an initial sequence element, V1 exhibits generative anticipatory responses even when the subsequent stimulus is withheld, strongly indicative of spatiotemporal pattern completion. However, our data also suggest that the brain may not store discrete elements of sequential experience in modular fashion, but rather as conjunctive representations of the stimuli as embedded within their spatiotemporal context (*58*). The well-controlled, passive visual stimulation protocols and robust electrophysiological reports of V1 plasticity described here provide a powerful platform to further delineate when and how the hippocampus contributes to neocortical encoding, storage, and retrieval of visual experiences.

## Materials and Methods

### Animal subjects

All subjects were male C57BL/6N mice obtained from Charles River Laboratory International (Wilmington, MA) at postnatal day 25-26 (P25-26). At the time that this study was conducted, only male mice were used due to evidence of distinct behavioral patterns in females, which are not adequately captured using our established assay of visually-evoked forepaw responses in head-restrained mice. This major limitation is being systematically addressed in the context of other studies. After arriving at MIT, mice were housed in groups of 2-5 littermates on a 12h dark-light cycle (light phase beginning at 7:00 a.m.) with food and water provided *ad libidum*. Nalgene homecages contained woodchip bedding and cotton nesting materials. All procedures adhered to the guidelines of the US National Institutes of Health and were approved by the Committee on Animal Care at MIT (Cambridge, MA, USA). All efforts were made to minimize pain or distress in the animals. Data reported is from experimentally-naïve animals except for the object displacement task, which provided an internal control for cohorts that previously underwent the electrophysiological recordings summarized in Figure 1.

### Surgical procedures

#### General surgical preparation

Surgeries were performed at P28-30 (except for the experiment in Fig. 5, which were performed at P35). Anesthesia was induced and maintained with inhaled isoflurane (1.5-3% in oxygen). Pre-operative subcutaneous injections of meloxicam (1 mg/kg) were administered as analgesic. Body temperature was maintained at 37°C with a heat source positioned under the torso of the mouse. Ophthalmic ointment was applied topically to both eyes to prevent damage. The scalp was shaved and cleaned with providine solution (10% w/v) followed by ethanol (70% v/v), and lidocaine hydrochloride (1%) was injected subcutaneously under the scalp as a local anesthetic. A 1 cm-long midline scalp incision was applied with surgical scissors to reveal an area of skull between the eyes and ears. To improve cement adhesion the skull surface was then carefully cleaned with saline, scored with a scalpel blade, and dried with compressed air. All mice went on to be implanted with local field electrodes. At the completion of all surgical procedures, mice were placed in a recovery chamber with free access to a heat source until the mouse regained consciousness and resumed grooming. In the lesion experiments, all mice received subcutaneous injections of warmed sterile Ringer’s solution to aid in recovery. Mice received daily subcutaneous meloxicam (1 mg/kg) injections as analgesic for 48-72 hours following surgery, and were monitored for signs of discomfort or illness.

#### Hippocampal excitotoxic lesions

The lesioning protocol was adapted from previous studies (*15*) through extensive pilot testing. Mice were prepared according to the general surgical procedures described above, then mounted on a stereotaxic apparatus (Kopf Instruments) with earbars. The pitch of the skull was adjusted such that bregma and lambda were level with the horizontal plane. A pulled glass pipette was backfilled with mineral oil and mounted on a nanoliter injector (Nanoject III, Drummond Scientific), then front-loaded with either freshly dissolved NMDA solution (10 mg/mL in sterile physiological saline [0.9% w/v]) in the Lesion condition, or an equal volume of sterile saline in the Sham control condition. Nano-infusions were targeted at 4 stereotaxically-determined sites per hemisphere, relative to bregma: 1) A/P - 1.8mm, M/L +/- 1.3mm, D/V (from dural surface above injection site) −1.4mm; 2) A/P −2.3mm, M/L +/-1.8mm, D/V (from dura) −1.6mm. To minimize the likelihood of damaging V1, for the two injections per hemisphere targeting ventral hippocampus the pipette was angled at 20 degrees, progressing ventrocaudally. Coordinates for these sites were adjusted to compensate for the angle of approach: 3) A/P −1.8mm, M/L +/- 2.9mm, D/V (from dura) −2.6mm; 4) A/P − 1.8mm, M/L +/- 2.9mm, D/V (from dura) 3.2mm. Tissue was permitted to decompress for 5 minutes after lowering the injector to the desired depth before commencing injections. At each of the first 3 sites 70 nL was injected per hemisphere, in 7 x 10 nL pulses at a rate of 43 nL/sec with 30 seconds between each pulse. At the fourth injection site 100 nL was delivered in 10 pulses per hemisphere. During infusions dura was kept moist by applying sterile saline to each craniotomy. The pipette was retracted 5-7 minutes following the final pulse, and the craniotomy was sealed with a small bead of Kwik-Sil silicone adhesive (WPI Inc.). Following the final infusion, diazepam was injected subcutaneously (5 mg/kg) as an anticonvulsant to reduce propagation of seizure activity during post-operative recovery after removal from isoflurane.

#### Electrode implantation

Extracellular local field potential (LFP) electrodes were implanted following the surgical procedures described above. A steel headpost was affixed between the eyes anterior to bregma using Krazy Glue (Elmers) followed by Loctite 454 cyanoacrylate adhesives. A dental drill was used to apply <0.5 mm craniotomies over frontal/motor cortex in the right hemisphere, and bilaterally over the binocular zone of V1 (+/3.1 mm lateral of lambda). A custom-fabricated silver wire (A-M systems, Sequim, WA, US) electrode was positioned approximately A/P +0.5mm, M/L + 1.0mm from bregma, and inserted to a depth D/V −0.3mm onto the surface of frontal cortex to serve as reference. Blunt tapered tungsten microelectrodes (300-500 MΩ, 75μm diameter; #30070, FHC, Bowdoinham, ME) were gradually advanced into binocular V1 to a depth of 450-470 um from dura, targeting layer 4. Electrodes were secured with cyanoacrylate glue, followed by application of Ortho-Jet dental acrylic (Lang Dental, Wheeling, IL, USA) to adhere to the skull surface. An adhesive accelerant (Zip-Kicker, Pacer Technologies, Ontario, CA, USA) was applied sparingly to expedite curing of cyanoacrylates.

### Visual stimulation

#### Grating stimuli

During all experimentation the investigators remained blind to group assignment. Visual stimulation and electrophysiological recordings were performed in one of two adjacent enclosed rooms, which remained dimly-lit throughout while preparing each animal and throughout experimentation. To avoid disruption of the circadian cycle by daily visual stimulation, experiments were performed during the 12-hour light phase. Awake, head-fixed mice were positioned with eyes 20 cm from the midline of a gamma-corrected LCD display. In one recording suite, the head fixation apparatus and monitor were both contained in a custom sound- and light-attenuating chamber coated on all external sides with grounded copper to minimize electrical noise. Internal walls of the chamber were matt black to reduce light reflections. In the second suite, the equipment was not enclosed within a chamber, but was well grounded away from sources of light, sound, and electrical interference. In both rooms, custom MATLAB software built around the PsychToolbox extension was used to precisely display fullfield visual stimuli. After recovery from surgery, mice were acclimated to head restraint in front of a 50% gray screen for 30 minutes on each of 2 days. The following day a binocular visual stimulation protocol was initiated, consisting of exposure to isoluminant sinusoidal grating stimuli (100% contrast, 0.05 cycles per degree) arranged in phase-reversing and/or sequential patterns (described below). The orientation of each stimulus was offset by at least 15° from the cardinal angles and 30° from all other stimuli: 15°, 45°, 75°, 105°, 135°, and 165°. Mice were placed in the head restraint apparatus 5 minutes prior to the beginning of each recording session, during which time a gray screen was presented.

#### SRP/orientation-selective behavioral habituation protocol

Mice viewed an oriented grating that phase reversed every 0.5 s (2 Hz). Daily for 6 days, mice viewed five blocks of 200 phase reversals interleaved by 30-sec presentations of isoluminant gray screen stimuli. During a SRP testing session (day 7), mice viewed gratings at both the familiar orientation and also a novel orientation (non-cardinal and 90° offset from previously-presented stimuli). Five blocks of each orientation were presented in pseudorandom order (no stimulus was presented for more than two consecutive blocks). A majority of mice reported in Figure 1 were also retested on day 17, during which the familiar stimulus (X°) was presented along with a second novel orientation.

In the static grating experiment, all mice viewed 6 x 100-sec blocks of a single grating orientation on days 1-6 of SRP induction, presented as static images of each contrast-inverted phase (3 blocks of each). On the 7^th^ day, two distinct orientations were each presented in a 2 Hz phase reversing pattern, including a novel orientation and the familiar orientations viewed over prior days. Both were presented in 3 interleaved blocks of 200 phase reversals.

#### Visual sequence protocol

On days 1-4 of the visual sequence protocol, a contiguous series of 4 oriented grating stimuli was repeated 200 times per day in a consistent spatiotemporal order. Individual sequence presentations were interleaved by 1.5-sec intervals of isoluminant gray screen, and arranged into 4 blocks of 50 repetitions interleaved by 30-second gray screen intervals. On test day (day 5), mice viewed 4 blocks of the same sequence in both forward and backward arrangements, as well as the forward sequence presented at 50% temporal frequency (each stimulus held for 300 ms, 1.5s interleaved gray screen). The order of blocks was pseudorandomly shuffled, such that no sequence was viewed more than twice consecutively. Another test session was presented 5-7 days later (experimental day 10), during which the familiar ABCD pattern was presented at the standard temporal frequency, as well as the same sequence with stimulus B omitted (A_CD). Only the first two stimuli (A and B) in the 50% temporal frequency and A_CD omission conditions were analyzed here. The specific grating orientation assigned to each position in the sequence was counterbalanced across mice.

#### Integrated SRP/Sequence protocol

To align the stimulation parameters used in the SRP and sequence protocols, in a modified paradigm individual mice were presented with two pairs of stimuli: one pair phase-inverted gratings of the same orientation (‘Flip’ and ‘Flop’), and the other two distinct grating orientations (‘A’ and ‘B’). The other parameters were adopted from the visual sequence protocol. Specifically, each stimulus was held on screen for 150 ms, with 1.5 sec gray screen stimuli presented between each pair. Four blocks of each stimulus pair were presented in pseudorandom order, with 30-sec gray screen periods interleaved. Only the plasticity induction sessions (Days 1-4) are reported.

### Electrophysiological recordings

Two electrophysiology systems (Recorder-64, Plexon Inc., Dallas, TX, US) were used to record neural LFP activity in awake, head-restrained mice throughout the multi-day visual stimulation protocols described above. Although these systems are comparable, the absolute magnitude of evoked responses recorded on each cannot be directly compared due to differences in amplifier configuration. Nevertheless, relative differences on each system can be compared (i.e. VEP potentiation, relative magnitudes of distinct VEP components, etc). Continuous extracellular voltage signals were collected bilaterally from V1 (referenced to the frontal electrode), low-pass filtered at 500Hz, and digitized at 1000 Hz. All data were analyzed with custom code written in MATLAB and C++. To extract visually-evoked potentials (VEPs), the 300-500 ms interval following each stimulus presentation was extracted and averaged across all trials within a block, or in some cases for specific trials across multiple blocks. VEP magnitude was generally defined as the voltage difference between the first negative-going trough following stimulus onset (~40 ms latency), and the subsequent positive going peak (typically occurring at a latency of between 65-125 ms). In **Figure 4A-C**, the subsequent N2 and P2 VEP components were also extracted, and the trough-to-peak magnitude was normalized the magnitude of the preceding N1 to P1 response to stimulus A. An automated detection algorithm was used to measure magnitudes, but each VEP was manually inspected to ensure consistent identification across experimental conditions. The same procedure was used to measure VEP magnitudes in the sequence-exposure protocol, with care taken to ensure that the positive-going component did not overlap with the evoked period of the subsequent stimulus. In the visual sequence paradigm, VEPs were averaged across each of the four stimuli (“ABCD”) when comparing response changes over days. However, generally the first stimulus in the sequence shows minimal response modulation as a function of familiarity, and cues anticipatory potentiation only for subsequent stimuli. Thus, to assess the effects of lesions on long-term plasticity, for the day-5 test session we opted to compare only responses to the stimuli in positions 2 and 3 of each sequence (orientations B and C) normalized to the ‘baseline’ magnitude for each mouse recorded on Day 1. Likewise, in the integrated SRP/sequence protocol, only the responses to the 2^nd^ stimulus in each phase- or orientation-shifted pair were compared between conditions. The second sequence element was also specifically measured in the temporal delay (ABCD_300ms_) and stimulus omission (A_CD) experiments (**Fig. 4**). Although electrodes were implanted bilaterally, recordings from only one hemisphere of each mouse were included in the final datasets. For each mouse, the chosen hemisphere was selected on the basis of the mean VEP with the largest mean magnitude on Day-1, and possessing morphology consistent with V1 layer 4. Following experimentation, electrode positions were confirmed histologically and those clearly falling outside of binocular V1 layer 4 were excluded from analysis (described below). All recordings were performed blind to group assignment.

### Visually-evoked behavior in head-restrained mice

In the head-fixed mice described above, behavioral data was obtained in parallel to all electrophysiological recordings. As we have not previously observed spontaneous behavioral responses selective for specific visual sequences, behavior was analyzed only during the SRP protocol. Mice were positioned with their forepaws on a modified piezoelectric pressure sensor (C.B. Gitty, #50-004-02) affixed directly beneath the head restraint bar, with the edge of the sensor resting inside the tube containing the animal’s torso (see **Fig. 1D**). The continuous analog signal was amplified, digitized, and recorded concurrently with the electrophysiological data using the Plexon Recorder-64 system. To quantify visually-evoked fidgeting (“vidget”) behaviors, the 1000 Hz voltage recordings were downsampled to 100 Hz, rectified (by subtracting mean voltage), and converted to absolute values. To obtain the average vidget response to the onset of each block of grating stimuli, the 5-sec of data collected immediately after onset (typically the first 10 phase reversals) were normalized to the mean activity of the preceding 10-sec of gray screen exposure. These normalized 5-sec intervals were then averaged across blocks and the mean value was computed to generate a vidget magnitude for each stimulus in a recording session (in arbitrary units; a.u.). To quantify spontaneous behavioral movement during gray screen (**Fig. S2A**) we simply averaged all 10-sec pre-stimulus intervals within a session for each mouse.

### Object displacement task

After completion of electrophysiological recordings, mice used in the SRP experiments were subjected to an object displacement task as a positive behavioral control to functionally confirm hippocampal ablation (*27, 28*). Mice were transported daily to a dimly lit room and permitted to acclimate for at least one hour prior to experimentation. Mice were then placed into a square open-field arena (40 x 40 x 30 cm) with clear plexiglass sides, located in the center of a large isolation chamber (as described above). Computer monitors were positioned on two opposing sides of the arena, each displaying isoluminant gray static stimuli to provide ambient lighting. Solid white two-dimensional geometric shapes were affixed to the black walls of the isolation chamber to serve as distal spatial cues that were each visible from any location within the arena. The first day of the task consisted of two 15-minute habituation sessions to an empty arena. On each of the next two days each mouse was returned to the arena for two 10-minute sampling sessions, during which the animal could freely explore two identical objects (small glass bottles filled with odorless colored liquid) positioned in opposite corners of the arena (approximately 5 cm away from the nearest walls). The position of the objects remained consistent across days, but varied across mice. Approximately 4-6 hours separated the sampling sessions each day. Approximately 24±1 hours from the final sampling session, the mice were returned to the arena, at which time one of the two objects had been shifted to a new corner.

Exploration was manually scored by an experimenter blind to treatment condition and object identity (i.e. familiar or novel position). Exploration was quantified as the duration of time the mouse spent actively investigating each object (mouse nose within 2 cm and facing towards the item while actively whisking). An *exploration ratio* was calculated by subtracting the time spent investigating the stimulus in the familiar position from the time spent investigating the displaced object (novel location), divided by total object exploration time. As mice rapidly habituate to new objects and their spatial positions, only the first 5 minutes of exploration was analyzed for each session.

### Post-mortem histology

To histologically quantify the extent of hippocampal lesioning, all mice were deeply anesthetized using Fatal-Plus (pentobarbital) and slowly perfused with 0.01M phosphate-buffered saline (PBS) followed by cold 4% paraformaldehyde (PFA). Brains remained in 4% PFA at 4°C for 24 hours before being transferred to PBS for storage. Fixed brains were sliced into 50 um sections using a vibratome and briefly stored in PBS-filled wellplates at 4°C. Every sixth slice (~300 μm increments) was then processed for NeuN neuronal nuclei immunohistochemistry to aid in visualizing hippocampal ablation (*59*).

For the immunohistochemistry procedure, multi-well plates containing floating slices were placed on a rotary shaker and incubated in blocking solution (20% fetal bovine serum, 0.2% Triton X-100 in 0.01M PBS) for 1 hour at room temperature. Following thorough aspiration of the blocking solution, slices were incubated overnight at 4°C in mouse anti-NeuN primary antibody (#MAB377, RRID:AB_2298772, Millipore Sigma, Billerica, MA) at a 1:1000 concentration in diluted blocking solution (10% fetal plus, 0.1% Triton X-100 in 0.01M PBS). After removing primary antibody, slices were washed 3 times in PBS and incubated for a further hour at room temperature in diluted blocking solution containing 1:500 goat anti-mouse Alexa488-conjugated IgG secondary antibody (#A28175, RRID:AB_2536161, Invitrogen, Carlsbad, CA) and 1:5000 Hoechst stain (Thermo Scientific, #33342). After three additional washes slices were mounted on charged glass slides (Superfront Plus, Fisher Scientific), briefly air-dried, then coverslipped with #1.5 glass using Prolong Diamond antifade mountant (Molecular Probes, #P36961). Slices were imaged with 2x and/or 4x objective lenses on a confocal fluorescence microscope (Olympus, Japan) and tiled using the FIJI distribution of ImageJ software (NIH).

For post-mortem histological verification of electrode placements (in non-lesioned cohorts), mice were deeply anesthetized via isoflurane inhalation and decapitated. The brain was carefully extracted and placed in 4% PFA for 48-72 hours at 4°C, then rinsed and transferred to 0.01M PBS for storage. Brain slices (50 um) were collected using a vibratome and mounted on charged glass slides, air-dried for approximately 24 hours, then processed using cresyl violet nissl stain. Slides were later coverslipped (#1.5 glass) with Permount mounting medium (Fisher Chemical, SP15). A confocal microscope was used to visualize Nissl staining using the bright-field channel. Positioning of electrode tracts (along with accompanying damaged) within V1 and hippocampus was assessed with reference to the mouse brain atlas (*60*).

### Data analysis and statistics

Throughout the results section, all data is expressed as group mean ± standard error of the mean (S.E.M.). Each dataset was assessed for normality and homogeneity of variance prior to choosing a statistical approach, using Levene’s, D’Agostino and Pearson, and, in the case of small sample sizes, Shapiro-Wilk tests. Two-way repeated-measures (RM) analysis-of-variance (ANOVA) were used to compare group responses across trials, blocks, days, or stimulus orientations. Greenhouse-Geisser was used to modify the degrees of freedom of repeated measures tests to correct for violations of sphericity. Interactions were followed by tests of simple main effects. Comparisons of individual values between groups (i.e. Day-1 VEP magnitudes or F/N ratios) were tested with unpaired, two-tailed t-tests. In some instances, differences between group means and baseline scores or F/N ratio parity were evaluated using one-sample *t*-tests. When datasets did not meet the assumptions required for two-way repeated-measures ANOVA, non-parametric methods were applied independently for each group (Friedman or paired Wilcoxon signed-rank tests). When appropriate, groups were then compared using ratios of responses to familiar and novel stimuli (Mann-Whitney *U* test). Contingent on the specific statistical test, Sidak’s, Dunnett’s, Dunn’s, or Bonferroni methods were applied to compensate for multiple comparisons. Uncorrected alpha was set to 0.05. Statistical analyses were performed with Prism 8 (GraphPad) and SPSS 25.0 (IBM).

#### Exclusion criteria

Animals were excluded on the basis of a number of predefined criteria. First, experimentation was discontinued in those exhibiting evidence of illness or distress, post-operative infection, eye damage, or behavioral abnormalities (i.e. excessive grooming, discoordination, etc). Mice were also immediately euthanized in the rare case of detachment of the electrode headcap. Mice were excluded from analysis if electrode placement fell outside of binocular V1 layer-4, as identified electrophysiologically and histologically. Furthermore, mice were excluded if mean VEP magnitudes during any session fell within 2 standard deviations of baseline “noise” (spontaneous potentials measured during gray screen). Among the cohort that received a 2^nd^ SRP test at Day 17, a subset of mice was not included. A littermate pair (1/group) was excluded due to declining LFP quality, and 2 mice/group were used to pilot a distinct protocol (not reported), and could not be retested. Exclusions were performed blind to group assignment, with the exception that each cage was required to contained at least one mouse from each experimental group. Although we did not use a paired design, if all cagemates from one group did not meet inclusion criteria then the littermates were also excluded.

## Acknowledgements

We are very grateful to Arnold Heynen, Suzanne Meagher, Nina Palisano, Jessica Buckey, Erik Sklar, Kiki Chu, Amanda Coronado, Athene Wilson-Glover, Katherine Marshall, and Erin Hickey for their instrumental administrative and technical support. We are also deeply indebted to Alyssa (Ying) Li for conducting pilot experiments, behavioral scoring, immunohistochemistry, and confocal histology, as well as Julie (Heejung) Kim, who assisted with histology. We would like to acknowledge Jeff Gavornik and Dustin Hayden for developing software used in the collection and analysis of this data. Thank you also to Sam Cooke and Ming-fai Fong for their scientific insights. This work was supported by the National Institutes of Health (R01EY023037), the Howard Hughes Medical Institute, the Picower Institute Innovation Fund, and the Picower Fellows program (PSBF).

## Supplementary figure legends

**Figure S1.**
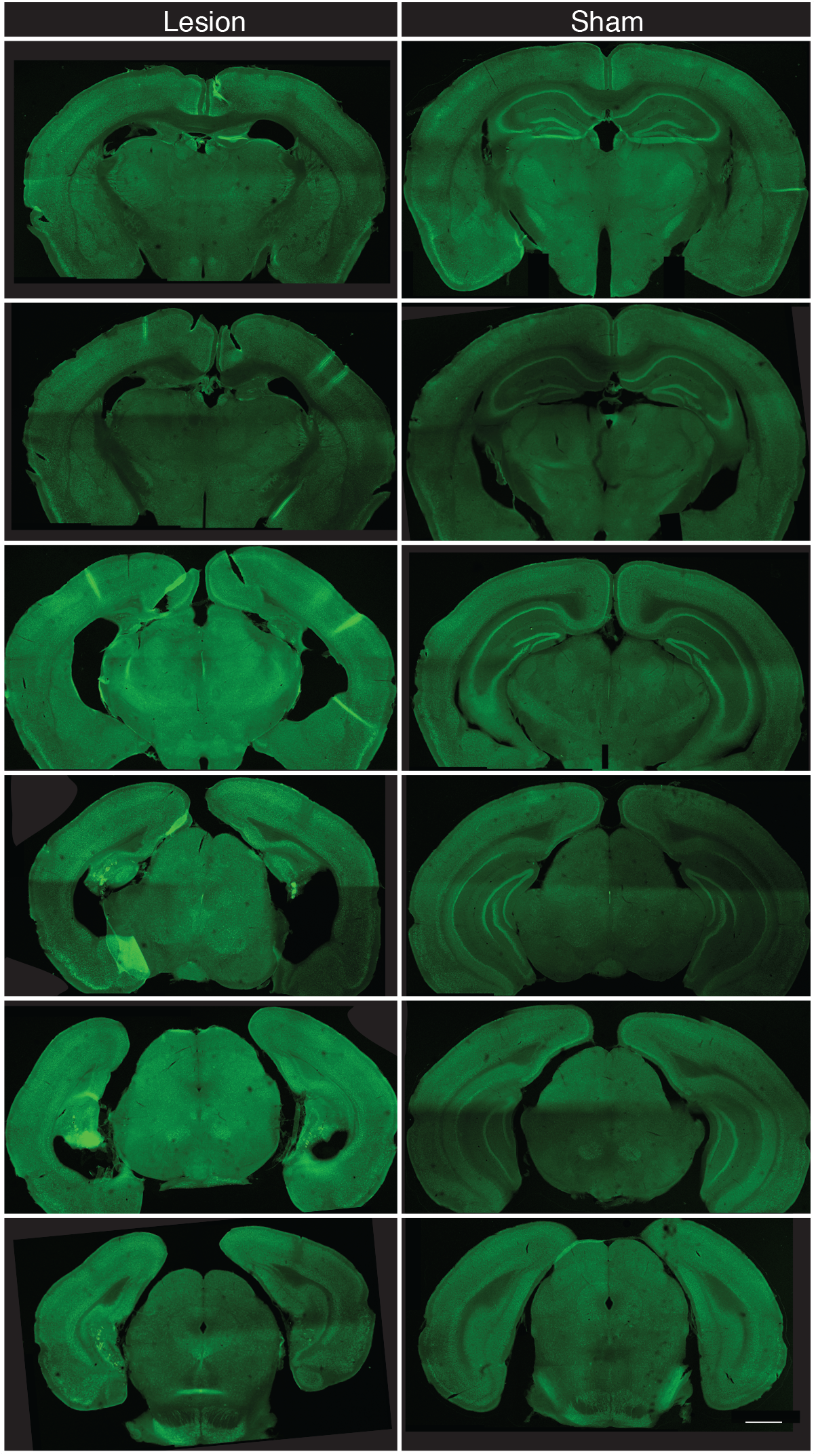
Hippocampal and sham lesion histology. Representative 50 μm coronal brain sections from hippocampal lesion (left panels) and sham control (right panels) mice. Green, NeuN; Blue, Hoechst stain. Scale bar represents 1000 μm.

**Figure S2.**
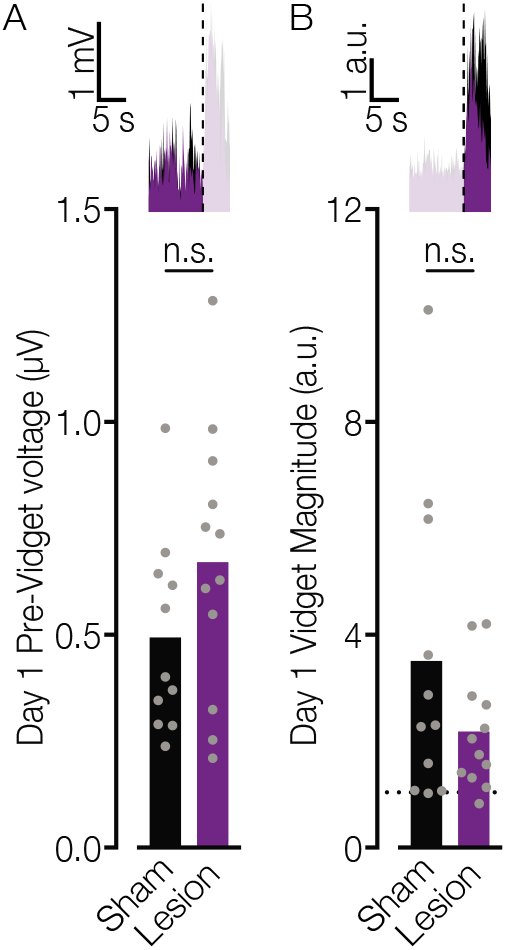
Spontaneous and stimulus-evoked behavioral responses on Day 1. **A)** Mean behavioral activity (in μV) during the 10 s baseline period prior to each stimulus onset is comparable for Sham and Lesion groups (mean voltage traces superimposed at top; unpaired *t*-test, *t*_21_ = 1.53, p = 0.14). **B)** Mean behavioral response magnitudes (arbitrary units, a.u.) elicited by visual stimulus onsets on *Day 1* also do not differ between groups (mean vidget traces superimposed at top; unpaired *Mann-Whitney U* = 52, *p* = 0.41). Both groups exhibit clear evidence of visually-evoked behavioral responses (one-sample *Wilcoxon* tests, Sham: *Z* = 2.845, *p* = 0.004; Lesion: *Z* =2.903, *p* = 0.004). Although not statistically significant, moderately elevated spontaneous baseline behavior is indicative of mild hyperactivity in the Lesion group (*61*), and could partially account for the slight reduction in their stimulus-evoked movement (which is normalized to pre-stimulus baseline).

**Figure S3.**
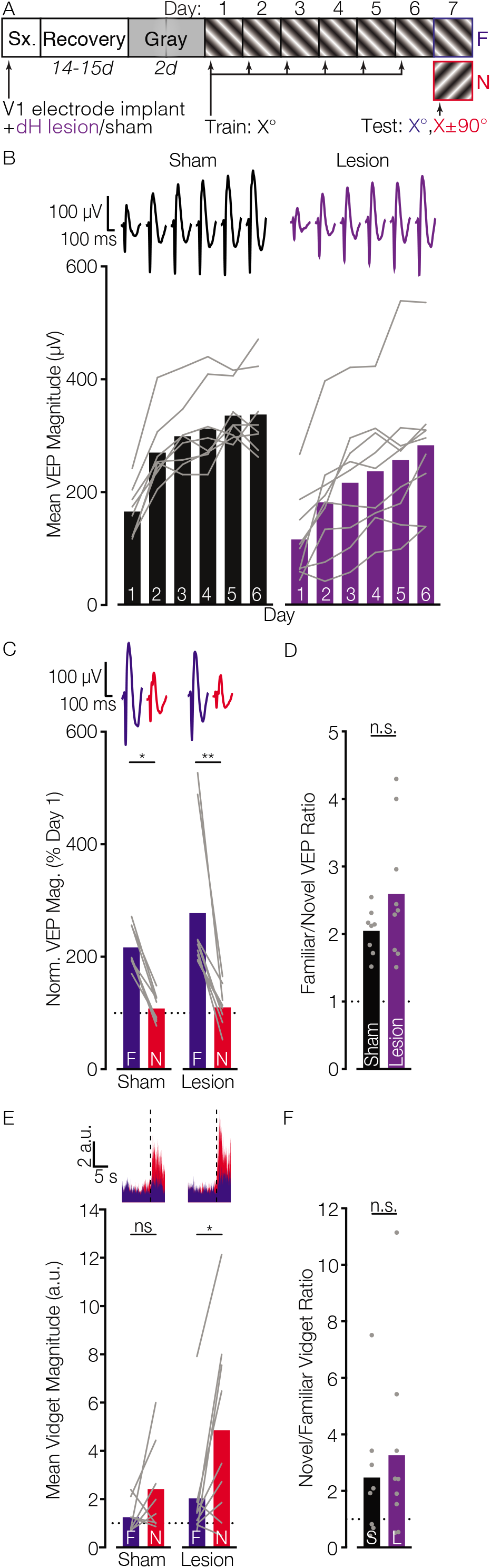
SRP and OSH are unaffected by lesions restricted to the dorsal hippocampus. **A)** Schematic summarizing the experimental protocol. **B)** Mice with dH lesions and sham controls exhibited significant VEP potentiation across days of exposure to an oriented grating stimulus (two-way RM ANOVA with Greenhouse-Geisser correction, *F*_3.242,48.63_ = 64.268, *p* < 0.0001; Sidak’s comparisons of *Day 1* vs *Days 2-6*, all p < 0.0001), and these groups did not differ statistically (main effect of group, *F*_1,15_ = 3.065, *p* = 0.1; group by day interaction, *F*_3.242,48.63_ = 1.046, *p* = 0.384). **C)** On *Day 7*, baseline-normalized VEP magnitudes in both the Sham and Lesion groups were larger for the F than N stimulus (two-tailed paired Wilcoxon signed-rank tests: Sham, *Z* = 2.52, *p* = 0.012; Lesion, *Z* = 2.67, *p* = 0.008). **D)** The F/N VEP ratio for the two groups did not differ (Welch’s two-tailed *t*-test, *t*_10.01_ = 1.56, *p* = 0.15). **E)** Paired Wilcoxon tests confirmed that on *Day 7*, mice with dH lesions vidget more to the N than F stimulus (*Z* = 2.31, *p* = 0.021), but there was no statistical difference in the Sham group (*Z* = 1.4, *p* = 0.16), in part due to small sample size. **F)** Nevertheless, the *Day 7* N/F vidget ratios for these groups were statistically equivalent (two-tailed Mann-Whitney *U* = 39, *p* = 0.85). *n*/group: Sham = 8, Lesion = 9.

**Figure S4.**
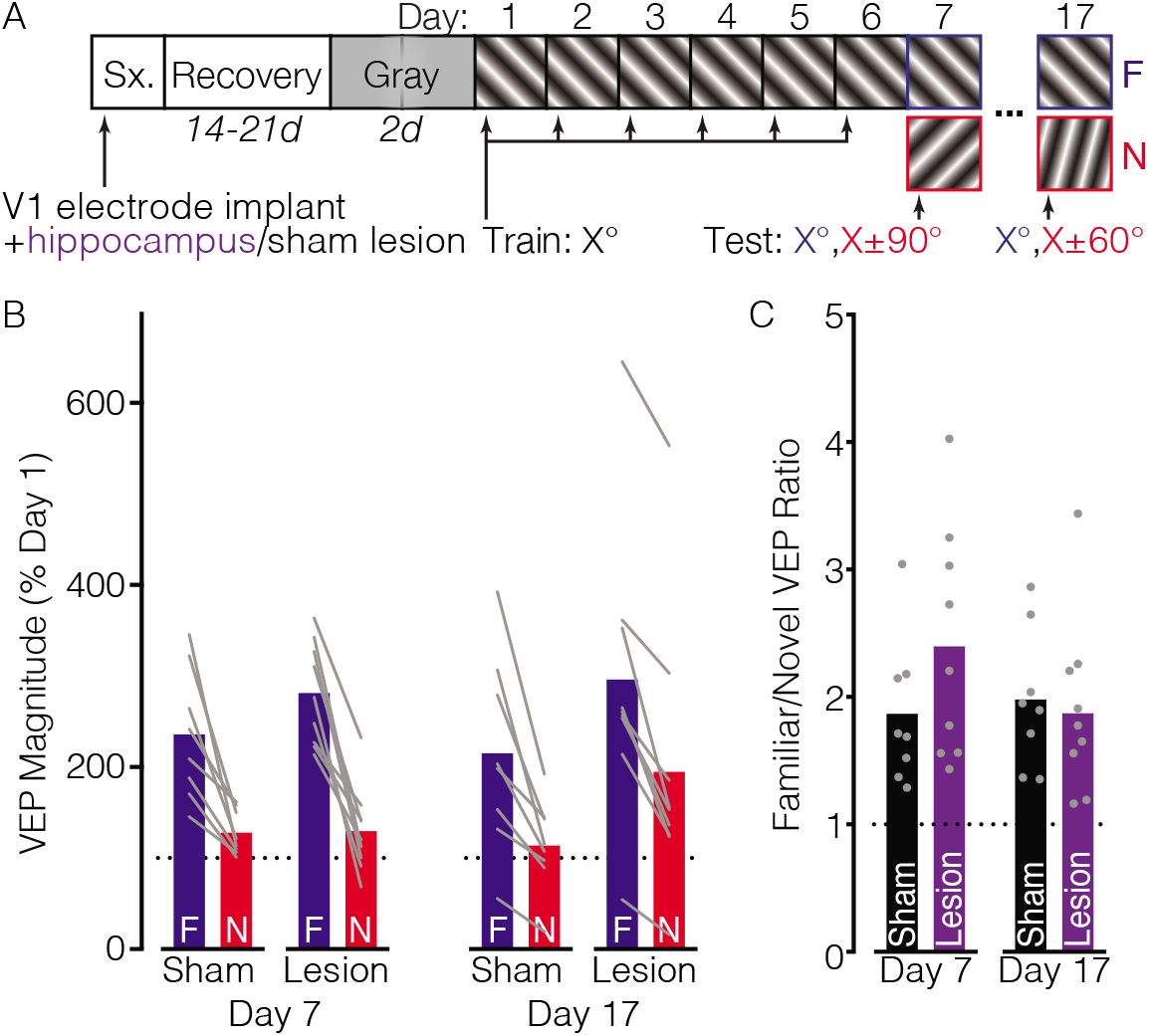
Impact of hippocampal ablation on SRP retention across prolonged intervals. **A)** A subset of mice reported in Figure 1 were retested 10 days following completion of the 7-day SRP protocol. Diagram summarizes the experimental timeline. **B)** Graphs plot *Day-7* and *Day-17* VEP magnitudes (normalized to *Day-1* baselines), and include only the subset of mice tested during both testing sessions (*n*/group: Sham = 8; Lesion = 9). As previously described, SRP expression was robust in both groups on Day 7 (two-way RM ANOVA, main effect of stimulus, *F*_1,15_ = 67.84, *p* < 0.0001). SRP also persisted when tested on Day 17 (*F*_1,15_ = 52.70, *p* < 0.0001). On Day 7 the normalized VEP responses elicited by the F stimulus appear exaggerated in the Lesion group (as reported in Fig. 1), but in this smaller cohort it does not reach statistical significance (no stimulus by group interaction, *F*_1,15_ = 1.91, *p* = 0.19). Similarly, on Day 17 the responses to both F and N appear mildly potentiated, although not a significant group difference (two-way RM ANOVA, no main effect of group, *F*_1,15_ = 1.80, *p* = 0.20; or stimulus by group interaction, *F*_1,15_ = 0.0002, *p* = 0.99). **C)** F/N VEP magnitude ratios during each testing session demonstrate that the slight SRP enhancement on Day 7 reaches the level on Sham controls by Day 17 (two-way RM ANOVA, no stimulus by group interaction, *F*_1,15_ = 4.27, *p* = 0.057).

**Figure S5.**
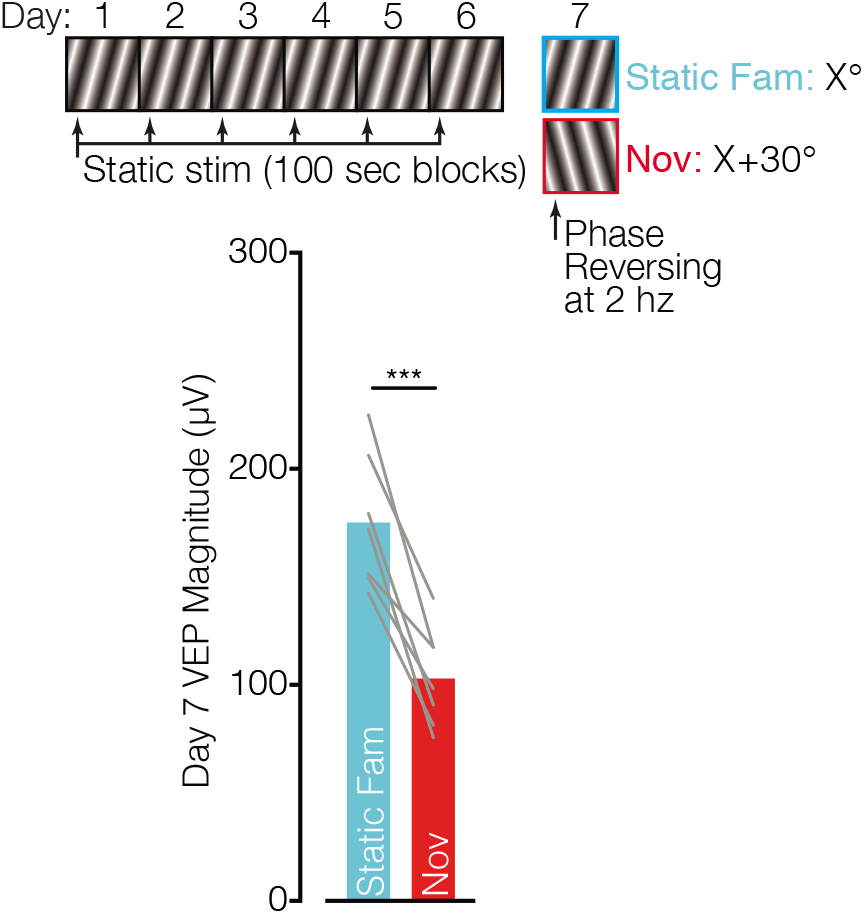
Daily exposure to static gratings elicits robust VEP potentiation. Diagram summarizes a modified SRP protocol in which each mouse (N = 7) viewed 6 x 100 sec blocks of a static grating stimulus during each of 6 daily recording sessions. On the 7^th^ day of the protocol all stimuli were presented phase reversing at 2 Hz, including a novel stimulus orientation along with the previously static stimulus (Static Fam). Static stimulation elicited robust SRP, revealed by comparison to the novel orientation (paired two-tailed *t*-test, *t*_6_ = 7.235, *p* = 0.0004). Thus, SRP does not require modulation of upcoming responses based on a predicted spatiotemporal pattern. *** < 0.001.

**Figure S6.**
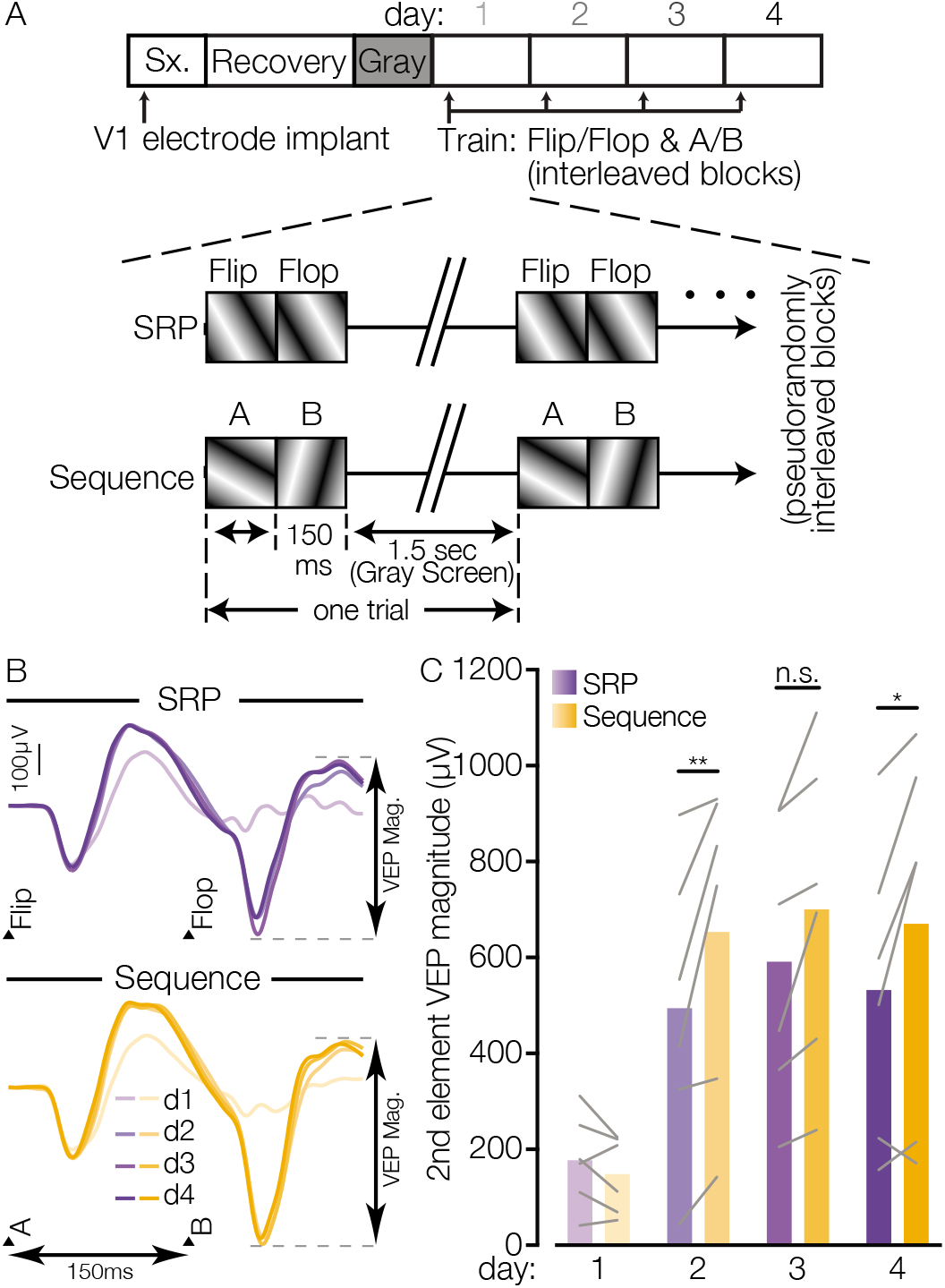
Orientation-shifted stimulus pairs elicit exaggerated potentiation compared to phase-shifted pairs. **A)** Diagram of a modified visual stimulation protocol combining the SRP and sequence protocols. Each mouse (N = 6) views two pairs of stimuli across interleaved blocks. The pairs of stimuli are either phase- or orientation-shifted (labeled ‘SRP’ and ‘Sequence’, respectively). All other stimulation properties are identical across the two conditions. **B)** Average VEP waveforms for the SRP and sequence stimulus pairs, with ticks denoting the onset of phase reversed (flip and flop) and orientation-shifted (A and B) images. **C)** Comparing VEP magnitudes elicited by the second stimulus in each pair (‘flop’ vs. ‘B’) indicates that potentiation over days is exaggerated for the orientation-shifted compared to phase-shifted stimulus (two-way RM ANOVA, Stimulus by Day interaction, *F*_3,15_ = 4.81, *p* = 0.015; Sidak’s posthoc comparisons of SRP and sequence VEPs on d1, *p* = 0.92; d2, *p* = 0.0036; d3, *p* = 0.050; d4, *p* = 0.011). We conclude that in addition to potentiation driven by familiarity with the identity of each oriented grating, during familiar visual sequences the brain predictively modulates responses to each cued stimulus, further enhancing VEP magnitude. * < 0.05, ** < 0.01, n.s. non-significant.

